# An anti-cancer binary system activated by bacteriophage HK022 Integrase

**DOI:** 10.1101/147736

**Authors:** Amer Elias, Itay Spector, Natasha Gritsenko, Yael Zilberstein, Rena Gorovits, Gali Prag, Mikhail Kolot

## Abstract

Cancer gene therapy is a great promising tool for cancer therapeutics due to the specific targeting based on the cancerous gene expression background. Binary systems based on site-specific recombination are one of the most effective potential approaches for cancer gene therapy. In these systems, a cancer specific promoter expresses a site-specific recombinase/integrase that in turn controls the expression of a toxin gene. In the current study, we have developed a new HK022 bacteriophage Integrase (Int) based binary system activating a Diphtheria toxin (DTA) gene expression specifically in cancer cells. We have demonstrated the efficiency, and the high specificity of the system *in vitro* in cell cultures and *in vivo* in a lung cancer mouse model. Strikingly, different apoptotic and anti-apoptotic factors demonstrated a remarkable efficacy killing capability of the Int-based binary system compared to the conventional *hTERT-DTA* mono system in the LLC-Kat lung cancer mice model; we observed that the active *hTERT* promoter down regulation by the transcription factors Mad-1 is the cornerstone of this phenomenon. The new Int-based binary system offers advantages over already known counterparts and may therefore be developed into a safer and efficient cancer treatment technology.

## INTRODUCTION

Cancer encompasses a large group of diseases characterized by the unregulated proliferation/apoptosis and metastasis, which represent one of the major healthcare problems. Thus, there is a strong unmet medical requirement for the development of novel therapies that provide improved clinical efficiency and longer survival time period in patients suffering with the different cancer types (1). For cancer therapies to be increasingly successful, however, the major obstacle that must be overcome is the safety efficacy of the cancer treatment technologies. Cancer therapy is approached by four main directions: *i*. Radiation therapy (XRT); *ii*. Chemotherapy; *iii*. Immunotherapy; and *iv*. Gene therapy (2). Despite that XRT is in common use it is associated with significant side effects on normal tissues and organs that limit the dosages and locations used (3). Similarly, use of the conventional cancer drugs in chemotherapy also has seriously side effects on the healthy cells, organs and whole organisms (4). The main reason of these harmful consequences is lack of tumor specificity. Moreover, the primary intrinsic and/or acquired multidrug resistance is one of the main obstacles to successful cancer treatment (5;6). Successful targeting of immune checkpoints to unleash anti-tumor T cell responses results in durable long-lasting response. However, such cancer immunotherapy helps only in a fraction of patients (7). Probably the combination of therapies is the most promising and effective approach in the cancer treatment (8).

Gene therapy is definitely one of the most important developing fields of the cancer treatment in since the early nineties, [one of the first studies was done by Zvi Ram (9)], and the technologies of many researches are currently in the advanced stages of clinical trials. The main advantage for gene therapy approach is the potential use of targeted delivery that mainly classified into two different strategies: passive targeting and active targeting; enhanced permeation and retention (EPR) is the basis of passive cancer targeting and which has been widely applied in numerous drug delivery systems for cancer targeting (10;11). Whereas, in active targeting, the basic concept is to utilize molecular targeting agent to specifically target the biomarkers or receptors on the cancer cells (12;13). Additional advantage of the gene therapy approach is the tumor-specific expression strategies to avoid harmfulness to normal tissue. However, the development of these tools still represents a significant challenge (2).

Toxin therapy approach as a part of cancer gene therapy is based on toxic gene expression targeted exclusively in the cancer cells resulting in killing through apoptosis without affecting healthy cells [as showed by Zvi ram in 1993, (9)]. One of the most used toxins for cancer gene therapy is the Diphtheria toxin (DT) of *Corynebacterium diphtheria* (14-16). DTA belongs to the A-B (Type III) group of exotoxins that catalyzes the ADP-ribosylation of elongation factor-2 (EF-2), hence arrests protein synthesis and induces cell death (17). DTA has a very efficient killing activity rate where a single DTA molecule is sufficient to kill the cell (18;19). Former studies demonstrated selective expression of DTA in tumor cells using a cancer specific promoter (5,17-21). The main drawback of this approach is the incomplete specificity of the promoter activity in cancer cells, whereas residual expression resulted in cytotoxic effect also in healthy cells (20;21).

Harnessing of site-specific recombinases (SSR) for genome manipulations and gene therapy is a well-known approach during the past 20 years (22-27). These systems encode recombinases that catalyze a site-specific recombination between two specific short DNA sequences of 30-40 base pairs (bp) that serve as recombination sites (RS) (28).

The site-specific Integrase (Int) of coliphage HK022 catalyzes phage integration and excision into and out of its *E. coli* host chromosome. The mechanism of these site-specific recombination reactions is identical to that of the well documented Int of coliphage Lambda (λ) (29). HK022’s host recombination site *attB* is 21 bp long and the phage *attP* recombination site is over 200 bp long. It is composed of a 21 bp core that is very similar to *attB* and flanked by two long arms (*P* and *P*’) that carry binding sites for Int and for some accessory DNA-bending proteins including the bacterial integration host factor (IHF). Integrative *attB* x *attP* recombination results in the integration of the phage into the host genome results in a prophage that is flanked by the recombinant *attL* and *attR* sites. The integration could be reversed in a process termed excision, which is also catalyzed by Int (30). We have demonstrated that Int of HK022 is active in human cells (31).

Site specific recombinases are also exploited in binary systems for the toxin specific expression in cancer cells that are based on two DNA plasmids. One is a recombinase substrate carries toxin gene silenced by a transcription terminator located between two Recombination sites, the second plasmid carries the recombinase gene expressed by a cancer specific promoter. This promoter provides the expression of the recombinase to the cancer cells where it excises the terminator and thus activates the toxin expression (9–11). We have shown previously that Int is much less active in eukaryotic cells compare to the Flp and Cre Site specific recombinases that are currently/ widely used for genome manipulations of higher organisms (32), and unpublished). This fact designates that the activation of a silent substrate of the binary system in case of Int demands considerably higher quantity of this enzyme that is reached only in cancer cells at the high level of activity of the cancer specific promoter. It is not the case in healthy cells where possible leakage of the cancer promoter may lead to basic level of Int expression that is obviously insufficient for such silent substrate activation. In case of much more active recombinase even insignificant level of such basic expression in healthy cells may lead to substrate activation and the toxin expression in the healthy cells. The described Int activity fine-tuning in the binary system insures the high efficacy safety level in the toxin based cancer therapy technology. We recently developed an Int based binary expression system and demonstrated its increased specificity in human cancer cells and in a lung cancer mouse model using luciferase as reporter (33). Here we demonstrate the application of the Int based binary system using a DTA toxin expression to specifically kill tumor cells for cancer therapy.

## MATERIALS AND METHODS

### Cells, growth conditions, mice, plasmids and oligomers

The bacterial host used was *E. coli* K12 strain TAP114 (*lacZ*) deltaM15 (34). It was grown and plated on Luria-Bertani (LB) rich medium with the appropriate antibiotics. Plasmid transformations were performed by electroporation (35). Plasmids and oligomers are listed in Tables 1 and 2, respectively.

**Table 1.** List of the plasmids

| Plasmid | Relevant genotype* | Source |
| --- | --- | --- |
| pEGFP-N1 | CMV-GFP, Km<sup>R</sup> | Clontech |
| pMK189 | CMV-attR-Stop-attL-GFP, Km<sup>R</sup> | (75) |
| pKS1161 | CMV-attR-Stop-attL-DTA, Km<sup>R</sup> | This work |
| pNA1263 | *hTERT*-Int-expressing plasmid, Ap<sup>R</sup> | (33) |
| pNG1805 | *hTERT*-DTA, Ap<sup>R</sup> | This work |
| pAE1808 | CMV-DTA, Km<sup>R</sup> | This work |
| pAE1850 | *hTERT*-attR-Stop-attL-DTA, Ap<sup>R</sup> | This work |

**Table 2.**
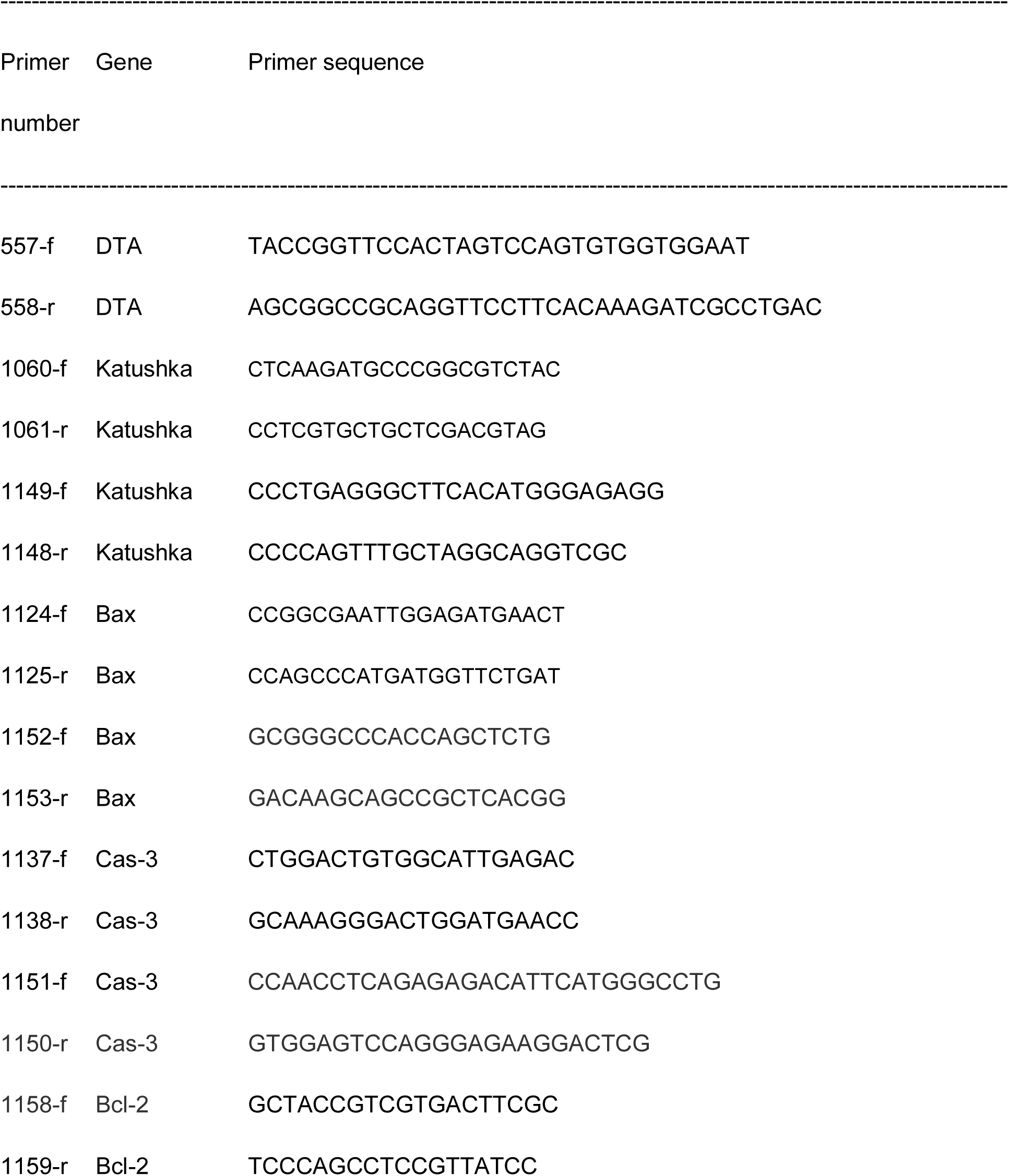
List of the primers

Human embryonic kidney cells HEK293 were cultured in Dulbecco’s modified Eagle’s medium (DMEM). Human normal skin fibroblast BJ cells were cultured in DMEM:M199 4:1 medium. Mouse Lewis lung carcinoma LLC1 cells labelled by Katushka fluorescent gene reporter (36) (LLC-Kat) were cultured in RPMI medium. For transient transfection of HEK293, LLC-Kat and BJ, the cells (~6x10^5^) were plated in a 6 well plate and 24 h later treated with 2.3 μg. circular plasmid DNA using CalFectin transfection reagent (SignaGen Laboratories, MD, USA) for HEK293 and TransIT-XT2 reagent (Mirus, WI, USA) for LLC-Kat and BJ.

All mice procedures were performed in compliance with Tel Aviv University guidelines and protocols approved by the Institutional Animal Care and Use Committee. C57BL/6 strain was used for the mice experiments.

All the experiments were at least repeated tri times.

### Plasmid construction

To construct DTA carried plasmids we used the less toxic DTA-176 mutant (G128D). (37). The plasmid pKS1161 carrying *CMV-attR*-Stop-*attL*-DTA cassette (Stop- transcription terminator) was constructed by a ligation of the AgeI-NotI DTA PCR fragment obtained using pIBI30-176 (kindly provided by Dr. Maxwell F.) as template and primers oEY557+oEY558 (Table 2, List of oligomers) into the same sites of pMK189. The plasmid pNG1805 carrying DTA gene under the control of *hTERT* promoter was constructed by ligation of the AgeI-blunted-NotI DTA fragment from pKS1161 with HindIII-blunted-NotI fragment of pNA1263. The plasmid pAE1808 carrying the DTA gene under the control of *CMV* promoter was constructed by a ligation of the AgeI-NotI DTA fragment from pKS1161 into the same sites of pEGFPN1. The plasmid pAE1850 used as a site-specific recombination substrate for DTA toxin assays under the control of *hTERT* promoter (*hTERT-attR*-Stop-*attL*-DTA) was constructed by a ligation of the NheI-blunted-AgeI *attR*-Stop-*attL* fragment from pMK189, AgeI-NotI DTA fragment from pAE1808 with HindIII-blunted-NotI fragment of pNA1263. Plasmid constructs were verified by DNA sequencing. (Table 1, List of plasmids).

### DTA toxin activity assay in cell cultures

For the estimation of Int dependent DTA toxin activity the appropriate cells culture (described above) were cotransfected by the GFP reporter plasmid (pEFGPN-1) (1 μg), site-specific recombination substrate plasmid pAE1850 (300 ng) and the Int expressing plasmid pNA1263 (1 μg). As a positive control of 100% GFP expression, the cells were transfected by plasmid pEFGPN-1 (1 μg). As a positive control of the DTA activity the cells were cotransfected by the plasmid pEFGPN-1 (1 μg) and the plasmids carrying DTA gene under the control of *CMV* (pAE1808) as non-tissue specific expression or *hTERT* promoter (pAE1805) as cancer specific expression (300 ng). As a negative control of Int dependent DTA toxin activity the cells were cotransfected by the reporter plasmid pEFGPN-1 (1 μg) and site-specific recombination substrate plasmid pAE1850 (300 ng). All cotransfection were performed in the total DNA quantity 2.3 μg. The mock plasmid DNA was pUC18. The efficiency of toxin activity was estimated by FACS analysis of the survival GFP expressing cells 48 hours post transfection.

### Fluorescent-activated cell sorting (FACS) analysis

~2 x 10^6^ cells from one well of a 6-well plate were collected following trypsin treatment of which 10^4^ cells were selected by the FACS sorter (Becton Dickinson Instrument) for fluorescent measurements of surviving cells. Data analysis was performed using the Flowing Software. Forward and side-scatter profiles were obtained from the same samples.

### DNA delivery in mice

DNA delivery in C57BL/6 mice was performed using the linear polyethylenimine based delivery reagent *in vivo*-jet PEI (Polyplus transfection, France) as previously described (33).

### Lung cancer development and fluorescence imaging in mice

0.8 × 10^6^ LLC-Kat cells were IV tail-injected into 8 week old (17–21 gr) female C57BL/6 mice. Lung tumor metastases development was examined nine and twelve days following the injection using *in vivo* micro-CT scanner (TomoScope^®^ *In-vivo* CT, Germany). The Katushka fluorescent signal in the lung metastases was analyzed on day 12 by an *in vivo* fluorescence imaging system (CRi MaestroTM, USA) (33).

### Preparation of sections, immunohistochemistry, confocal microscopy, apoptosis TUNEL assay and image analysis

Preparation of sections, confocal microscopy, apoptosis TUNEL assay and image analysis were performed as previously described (33). Healthy and cancer lungs carried metastases were immune-stained with anti-DTA (Cat#4701, ViroStat, Portland, ME, USA), cleaved Cas-3 (Cat#9664S, Cell Signaling, MA, USA), P-53 (Cat# SC-6243, Santa Cruz Biotechnology, USA), Mad1 (Cat# 4682, Cell Signaling, MA, USA), Katushka (Cat#AB233, Evrogen, Moscow, Russia) antibodies at 1:200 dilution. Donkey anti-Goat IgG (H+L) secondary antibodies conjugated with Cy2 dye (Cat# 705-225-147, Jackson ImmunoResearch, PA, USA) was used for DTA detection. Donkey anti-Rabbit IgG (H+L) secondary antibodies conjugated with Cy3 dye (Cat#711-165-152, Jackson ImmunoResearch, PA, USA) was used for cleaved Cas-3, P-53, Mad1 and Katushka detection.

### Immuno-detection of cellular protein extracts

Mouse lungs were dissected, washed in CMF buffer (137 mM NaCl, 2.7 mM KCl, 8 mM Na_2_HPO_4_, 1.5 mM KH_2_PO_4_, 5.5 mM glucose) and homogenized, 10 strokes of pestle A and 10 strokes of pestle B in a Dounce homogenizer, in lysis buffer [1 M sorbitol, 10 mm HEPES (pH 7.5), 5 mm EDTA, 0.25 M NaCl, 0.2 % Triton X-100, 0.2 % NP40 and complete protease inhibitor mixture (Roche Applied Science)]. After 30 min incubation on ice with vortex, the protein extracts were cleared by spin-down, and a standard PAGE loading buffer supplemented with 2% SDS was added. Samples were incubated at 65 ^o^C for 15 min and subjected to 12 % SDS PAGE. Immunodetection of cellular proteins was performed by western blotting according to standard procedures. Antibodies used for western blotting included anti-cleaved Cas-3, Katushka, anti-phospho-SAPK/JNK, anti-phospho-ERK (Cat#9251S and Cat#9101S, Cell Signaling Technology, USA) and anti-actin (Cat#ab180, Abcam, Cambridge, United Kingdom). Incubation with peroxidase-coupled secondary antibodies (Sigma–Aldrich, USA) was followed by the enhanced chemiluminescence detection procedure (Amersham, Bucks, UK).

### Quantitative Real-Time PCR

Total RNA was isolated from murine lungs with EZ-RNA Isolation kit (Biological Industries, Beit Haemek, Israel). cDNA was synthesized using Verso cDNA kit (Applied Biosystems, Foster City, CA). mRNA expression was assessed by StepOne quantitative Real-Time PCR system (Applied Biosystems, Foster City, CA). qRT-PCR for mouse Katushka, Cas-3, Bax and Bcl-2 mRNA were carried out. Details of the primers are given in Table 2.

In order to normalize the expression of Cas-3, Bax and Bcl-2 genes according to Katushka cells distribution in murine lungs, Absolute Quantification (AQ) method was applied. This method is based on standard curve of known quantity, with comparison of unknown quantities to the standard curve and extrapolation of the expression value. First, total number of mRNA molecules of Katushka, Cas-3, Bax and Bcl-2 were calculated according to its expression using the formula: μg DNA x pmol/660 pg x 10^6^ pg/1 μg x 1/N = pmol DNA. Then the number of Cas-3, Bax and Bcl-2 molecules was divided by the number of Katushka mRNA molecules in order to normalize their gene expression to one Katushka molecule.

### DNA manipulations

Plasmid DNA from *E. coli* was prepared using a DNA Spin Plasmid DNA purification Kit (Intron Biotechnology, Korea) or a NucleoBond^TM^ Xtra Maxi Plus EF kit (Macherey-Nagel, Germany). General genetic engineering experiments were performed as described by Sambrook and Russell (35).

### Statistical analysis

Data were presented as the mean ± SD. *P*-values of less than 0.05 were considered statistically significant by Student’s two-tailed t test assuming equal variance.

## RESULTS

### Integrase based binary DTA expression system for specific cancer cell killing

We previously demonstrated the validity of the Int activated cancer specific binary cell expression system with the *luc* reporter (33). Here we report the application and the level of specificity of the Int activated binary system using the DTA toxin to kill cancer cell (henceforth the binary system, Fig. 1A). This binary system consists of two plasmids: a pNA1263 that expresses the *int* gene under the control of the cancer specific *hTERT* promoter that is active practically in all types of tumors and immortal cells, but is silent in somatic tissues (38;39), and a second plasmid pAE1850 that carries the silent open reading frame of the *dta* gene separated from the *hTERT* promoter by a transcription terminator (40)(Stop) flanked by tandem *attR* and *attL* HK022 recombination sites (Fig. 1A). Int-catalyzed *attR* x *attL* recombination excises the terminator sequence, thereby allowing expression of the toxin from the promoter. In this binary system the expression of both genes (*int* and *dta*) are regulated by the *hTERT* promoter. As a positive control of the DTA toxicity in cell cultures we used plasmid pAE1808 that expresses *dta* gene from the constitutive and strong human cytomegalovirus promoter (*CMV*). To exanimate the efficiency of the binary system versus the conventional mono DTA-based cancer gene therapy approach; we used the plasmid pNG1805 that carried *dta* gene under the control of *hTERT* promoter.

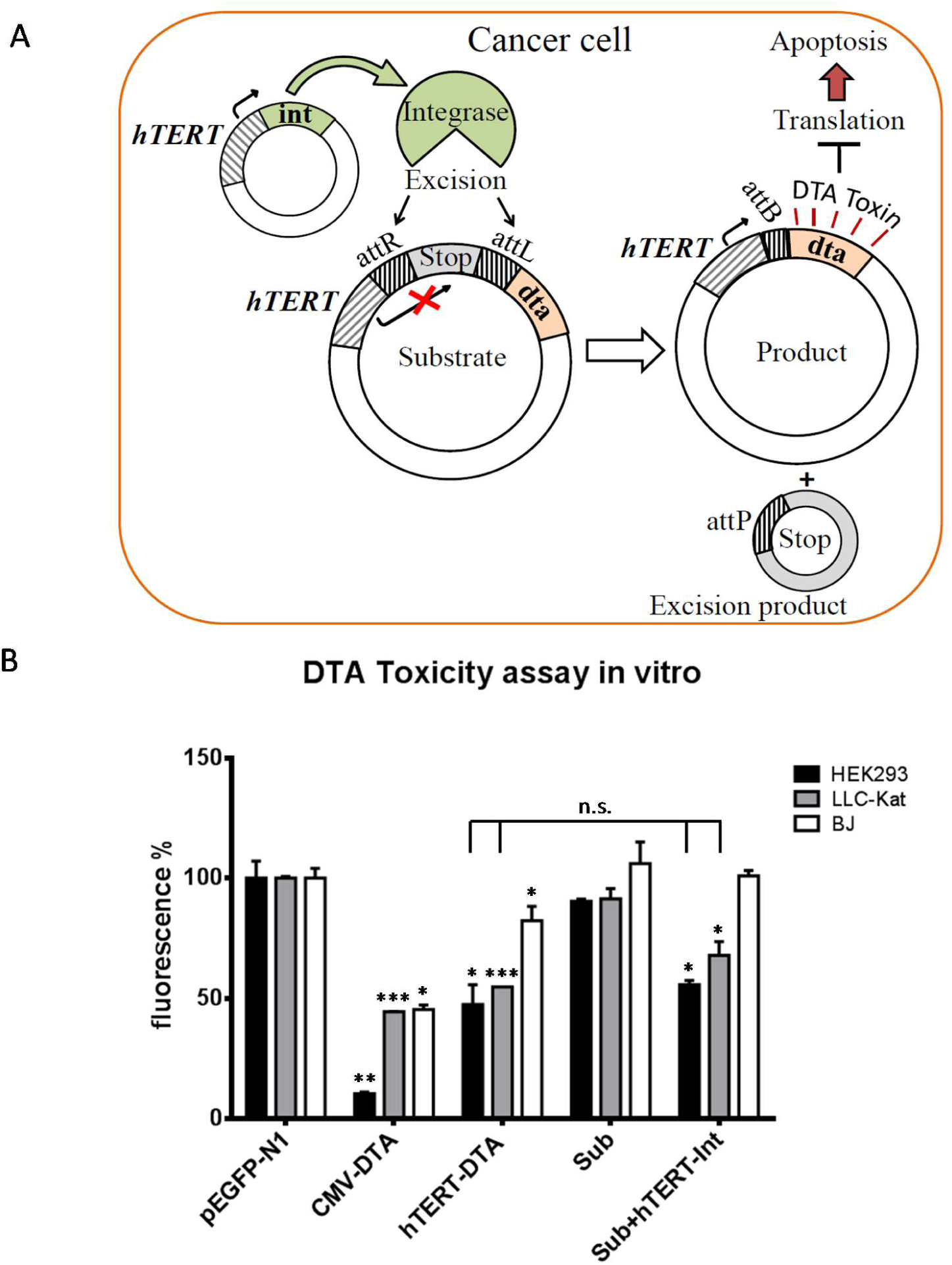
Figure 1. A. Scheme of the Int-based binary system. The recombination substrate carrying a silenced DTA toxin gene separated from its *hTERT* promoter by a transcription terminator (Stop) flanked by *attR* and *attL* recombination substrates. The recombination products generated by Int excision reaction: the activated DTA gene separated from its *hTERT* bacterial recombination site *attB* (top), and the excised circular attP-Stop (bottom). B. DTA toxicity assay *in vitro* in HEK293 and LLC-Kat (black and gray columns, respectively). C. DTA toxicity assay in vitro in BJ and A549 cells (white and gray columns, respectively). The cells were co-transfected with pEGFP-N1 (GFP expressing plasmid) and the following plasmids: CMV-DTA; *hTERT*-DTA; Sub; Sub + *hTERT*-Int, each compared with the EGFP-N1 only. Surviving cells tested by the estimation of GFP fluorescence level- Fluorescence % - percentage of the pEGFP-N1. The bars show the mean value of four experiments; the error bars indicate standard deviation. n.s: nonsignificant, ***p-val ≤ 0.001; **p-val ≤ 0.01; *p-val ≤ 0.05 vs. pEGFP-N1.

### The efficiency and specificity of the Int activated binary system in tissue cultures

First, we have examined the toxic activity of the less active dta mutant (G128D) based on its ability to abolish the translation of the green fluorescent protein (GFP) as reporter in cell cultures (37). The cell lines: HEK293, LLC-Kat and BJ were cotransfected with three plasmids: the silent dta substrate plasmid that carried the *hTERT* promoter-*attR*-Stop-*attL-dta* cassette (henceforth the sub), the *hTERT*-Int expressing plasmid and the EGFP-expressing plasmid (pEGFP-N1). The same cells were transfected with two control plasmids: one to verify the DTA activity using *CMV-dta* and the second carries the *hTERT- dta* expressing plasmid, each cotransfected with the EGFP reporter plasmid. To assess the essential silencing of the sub cells were cotransfected with sub alone and the pEGFP-N1 plasmids. The efficiency of the DTA activity was monitored by comparing EGFP expression in surviving cells 48 hours post transfection of each treatment compared to cells transfected by the EGFP plasmid alone. Three cell lines each underwent these five transfections: The first line (HEK293) consisted of immortalized human embryonic kidney cells (Fig. 1B black columns), that unlike normal human cells express a high level of *hTERT* sufficient to maintain their telomeres indefinitely (41;42). Therefore, *hTERT-int* expression of the binary system is expected to activate DTA toxicity of the sub plasmid. Quantitative FACS data in this cell line have demonstrated no toxicity in cells transfected with the silent sub alone (91% surviving cells that shows non-significant difference from the positive 100% control with the pEGFP alone) (Fig. 2, black column in Sub). The cells showed strong toxicity (10% surviving cells) with the *CMV-dta* plasmid and significant and comparable toxicity (47% and 55%) with the mono *hTERT-dta* and the binary systems (Fig. 1B, black columns). The second treated line (LLC-Kat) also known as Katushka cells, consisted of Lewis lung carcinoma line 1 cells (43) labelled with a far-red mutant of the red fluorescent gene of the sea anemone Entacmaea quadricolor (36;44) (Fig. 1B, grey columns). These cells were used in our previous report (33) to develop lung-metastatic tumors in mice (33). The data with this cell line also have demonstrated no toxicity in cells transfected with the silent sub alone (92% surviving cells that shows non-significant difference from the positive 100% control) (Fig. 1B, gray column in Sub). The cells showed strong toxicity (44% surviving cells) with the *CMV-dta* plasmid and significant toxicity (55% and 73%) with the mono *hTERT-dta* and the binary systems (Fig. 1B, gray columns). It should be noted that DTA activity in vitro in LLC-Kat cells was lower than in HEK293 cells (Fig. 1B, grey and black columns in *CMV-dta*).

**Figure 2.**
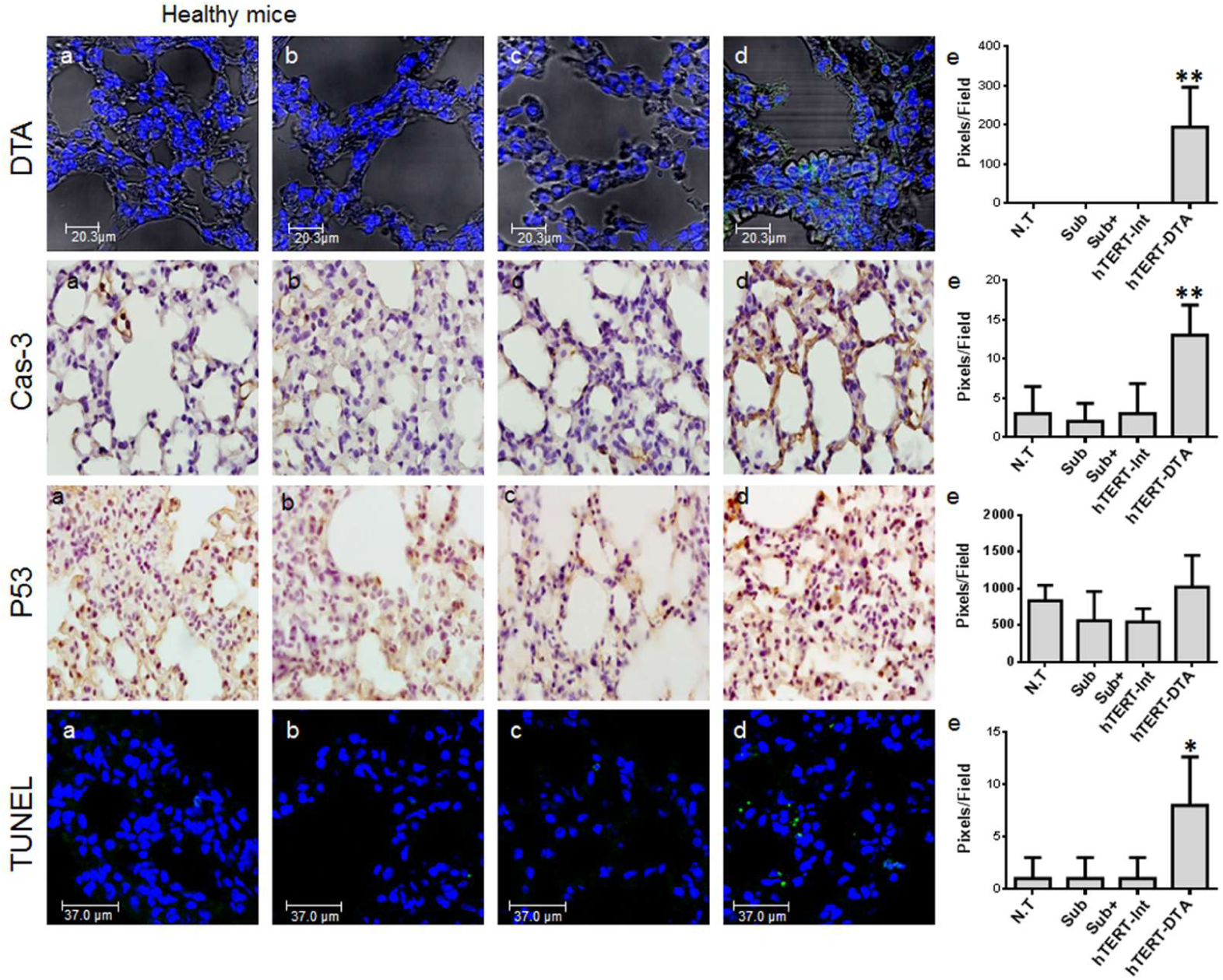
IHC analyses of healthy mice lungs: DTA toxin, activated Cas-3, P53 expression and TUNEL assay. (a) Untreated mice (N.T.); (b-d), IV-tail injected mice with the following plasmids complexed with the *in vivo-*jetPEI: (b) Sub (40mg), (c) Sub (40mg) + *hTERT*-Int (10mg), (d) *hTERT*-DTA (40mg), (e) Quantitative data. Each figure shows a typical lungs biopsy specimen from a cohort of at least five mice. The bars show the mean value of five experiments; the error bars indicate standard deviation. *p-val ≤ 0.05; **p-val ≤ 0.01 vs. N.T.

The third treated cells line (BJ) consisted of normal human foreskin fibroblast cells aimed to examine the cancer specific efficacy of the *hTERT* promoter (Fig. 1B, white columns). The data in this cell line also have showed no toxicity in cells transfected with the silent sub alone (100% surviving cells) (Fig. 1B, white column in Sub). The cells showed strong toxicity (45% surviving cells) with the *CMV-dta* plasmid and significant toxicity (82%) with the mono *hTERT-dta* (Fig. 1B, white columns). On the other hand, the binary system driven by *hTERT* promoter did not demonstrate any toxicity compared to the cells transfected by the pEGFP alone (Fig. 1B, white column in Sub+ *hTERT-int*). These results clearly showed that Int based DTA expression system is highly efficient to kill lung cancer cells in vitro while presented highly safety functionality in normal cells as the DTA expression was found to be essentially silence probably due to the transcription Stop sequence.

### Int-based binary system for DTA expression selectively kills mice cancer cells

At the next stage, we wanted to examine the cancer specific efficacy in killing as well as in safety of the Int-based binary system in whole organism *in vivo*. Mice were treated with the binary system and the therapeutic effect was assessed after 24 hours.

Four groups (6 mice/group) of two mouse types; healthy and LLC-Kat mice were treated with different plasmids combinations complexed with the *in vivo*-jet PEI reagent by IV tail-injection [panels (b-d) in Figs. 2-3]. Group (a) of mice in these figures remained untreated. Group (b) carried the sub alone to assess the absence of DTA leakage. Group (c) carried the sub and *hTERT-int* expressing plasmid to examine the efficacy of safety in healthy mice and cancer specific efficacy killing in lung cancer LLC-Kat mice of the binary system. Group (d) carried *hTERT- dta* expressing plasmid, as an example of the conventional DTA-based cancer gene therapy approach. 24 hours post injection; the biopsy specimens from all groups’ lungs were collected and used for the further analyses.

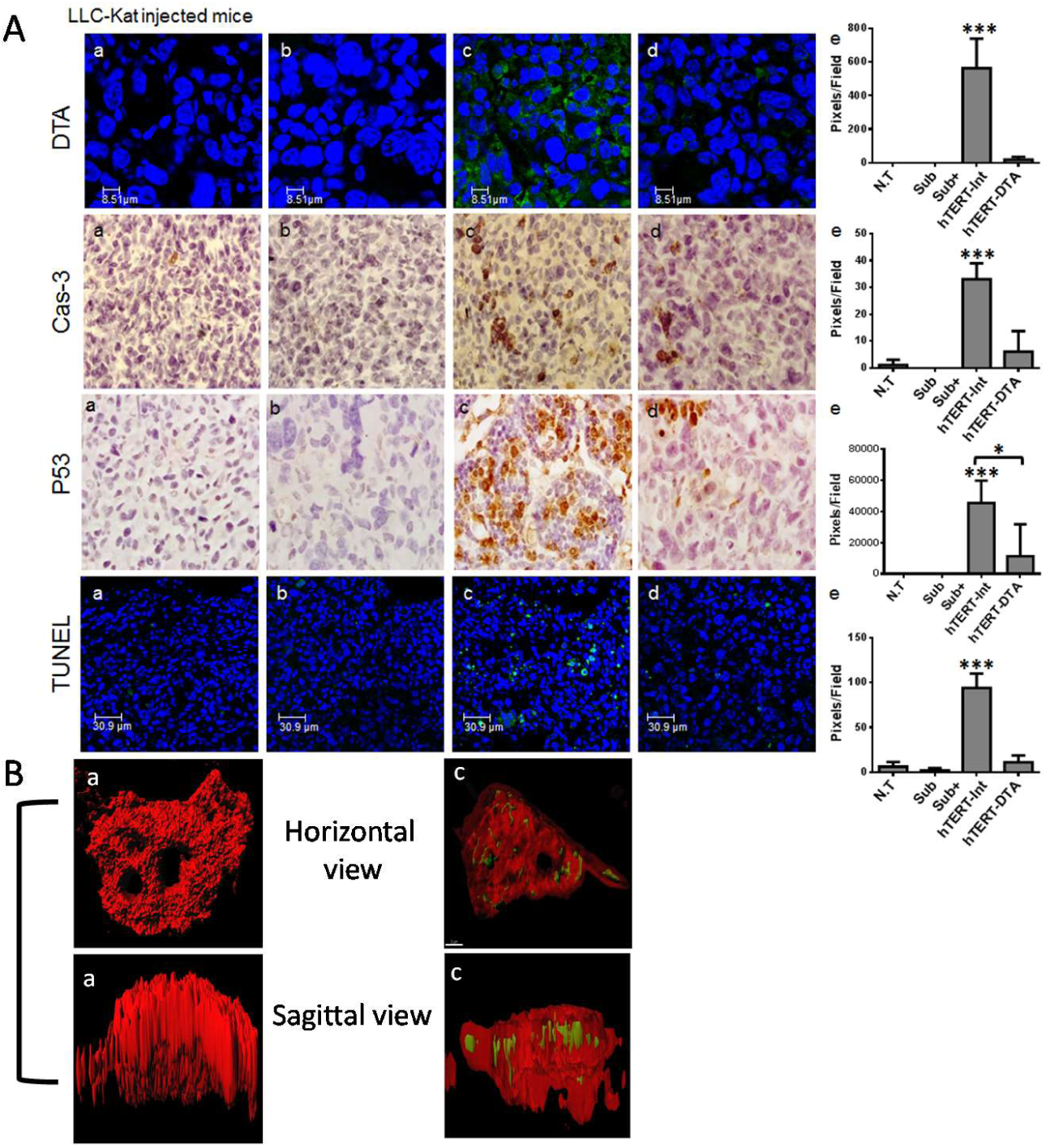
Figure 3. A. IHC analyses of LLC-Kat injected mice lungs: DTA toxin, activated Cas-3, P53 expression and TUNEL assay. The mice were treated in the same conditions as in Fig. 3. (a) Untreated mice (N.T.), (b) Sub, (c) Sub + *hTERT*-Int, (d) *hTERT*-DTA, (e) Quantitative data. Each figure shows a typical lungs biopsy specimen from a cohort of at least five mice. The bars show the mean value of five experiments; the error bars indicate standard deviation. *p-val ≤ 0.05 vs. *hTERT*-DTA; ***p-val ≤ 0.001 vs. N.T. B. Single cell 3D analysis of the DTA cancer specific expression (green) in the LLC-Kat cell (red) of the cancer mice lungs. The mice were IV-tail injected with the Sub (40mg) + *hTERT*-Int (10mg) plasmids complexed with the *in vivo-*jetPEI. Each figure shows a typical lung sample from a cohort of at least five mice. a- horizontal view, b- inside the cell, horizontal view, c- inside the cell, coronal view, d-sagittal view. Non-DTA expressing cell (e-f): e- horizontal view, f- sagittal view.

First, to demonstrate the cancer specific DTA expression in mouse lungs under the conditions described previously, an immunohistochemistry analysis (IHC) by DTA anti-bodies (DTA-Ab) was performed (Figs. 2 and 3). Among the mice treated with the Int-based binary system [Figs. 2 and 3, panels (c)], only the LLC-Kat mice showed substantial DTA positive immune detection [Fig. 3, panels (c, e)] suggesting that is selectively expressed and function in lung cancer cells *in vivo* in mice model. However, among the mice treated with *hTERT- dta* plasmid, positive DTA immune detection was observed both in cancer and normal mice [Figs. 2 and 3, panels (d, e)].

Comparative quantitative analysis of DTA expression level by IHC in LLC-Kat mice lungs has revealed 33 folds of DTA immune detection of the developed Int-based system over the *hTERT-dta* mono system (Fig. 3, e). We detected a substantial increased DTA expression in healthy mice that were treated with *hTERT-dta* plasmid compare with same mice that treated with the Int-based binary DTA expression system demonstrating the highly efficacy safety of the binary system (Fig. 2, e).

Since DTA expression induces apoptosis (18;19) in the following experiments the levels of apoptotic effect was analyzed in the lung specimens of healthy and LLC-Kat lung cancer mice using IHC, Western blot and quantitative Real-Time PCR (qRT PCR) methods.

First Caspase-3 (Cas-3) levels were detected by IHC (Figs. 2 and 3, Cas-3); Cas-3 is the most extensively studied apoptotic protein among caspase family members which expresses at the early stage of apoptosis (45). The immuno-detection of cleaved Cas-3 (at Asp175) distinguished endogenous levels of the large fragment of activated Cas-3, was clearly detected. Similarly, we detected significant elevation of P53 in the binary system treated mice compare with all other treated mice. P53 is the key regulator of cell cycle and well known to function as apoptotic marker (Figs. 2 and 3, P53) (46). Moreover, an associate significant elevation was also detected with the TUNEL assay (Figs. 2 and 3, TUNEL). TUNEL is a common method for identification of the final stage of apoptotic DNA fragmentation (47). Notably, the absence of apoptotic response was found in the lungs of healthy and LLC-Kat mice treated with the silent dta substrate, demonstrating the low level of apoptosis in these lungs. These results also indicate the absence of DTA leakage [Figs. 2 and 3, panels (b, e)]. Among mice treated with the Int-based binary system only the LLC-Kat ones showed the substantial immune detection by IHC for Cas-3, P-53 and TUNEL, suggesting high specificity of the binary system toward cancer cells [Fig. 3, panels (c, e)]. Unanticipated positive immune detection has been received in both types of mice treated with the *hTERT- dta* plasmid [Figs. 2 and 3, panels (d, e)].

Comparative quantitative analysis of the immune detected levels of the Cas-3, P53 and TUNEL in the lung specimens of LLC-Kat mice of Int-based binary and *hTERT-dta* mono systems has revealed six, five and nine folds respectively higher in the proposed binary system than the mono system [Fig. 3, panels (e)].

Finally, analysis of Int-based binary system for DTA expression has revealed its safety efficacy in the healthy mice [Fig. 2, panels (e)] along with significant killing activity in LLC-Kat lung cancer ones [Fig. 3, panels (e)] compare to the mono *hTERT- dta* system.

Having demonstrated the Int-based binary system killing activity of cancer cells *in vivo* as a result of selectivity expression of DTA in cancer cells we wanted to visualize the DTA expression in cancer mouse lungs by single cell analysis. IHC 3D analysis by double immunostaining using antibodies against Katushka and DTA (Fig. 3B, Red and Green respectively) of the LLC-Kat mice treated with the Int-based binary system presented significant amount of DTA in the cytoplasm of the single cancer cell demonstrating the high level expression of the DTA toxin in the cancer cells (Fig. 3B).

### The effect of the Int-based binary system on the apoptotic and tumor specific genes level expressions in cancer lung tissues

To further support the IHC results of the binary system in LLC-Kat cancer mouse lungs, we have performed a Western blot and qRT PCR analysis to reveal the effect of this system on different apoptotic, anti-apoptotic and tumor specific genes expression level. Western blot analysis was executed using the antibodies against apoptotic protein markers, such as cleaved Cas-3 and activated/phosphorylated Jun N-terminal kinase (JNK-Ph) (48;49), antibodies against proliferation marker: phosphorylated mitogen-activated protein kinase (ERK-Ph) (50;51) and the antibodies used as positive controls: Katushka and Actin. Cellular proteins were extracted from the lungs of an untreated and treated with the binary system LLC-Kat cancer mice (Fig. 4A, N.T. and Sub+*hTERT-int*, respectively). Fig. 4A shows that the Katushka patterns were detected in both cancer lung samples, confirming the cancer development. ERK-Ph proliferation marker abundance was decreased in the lungs of the mice treated with the binary system compare to untreated mice (Fig. 4A). On the other hand, both apoptotic markers Cas-3 and JNK-Ph showed essential increased abundances in the lungs of the mice treated with the binary system (Fig. 4A). The differential levels of the apoptotic and proliferation markers in LLC-Kat cancer mice treated with the Int-based system compared to the untreated mice presented the efficiency of the applied cancer treatment.

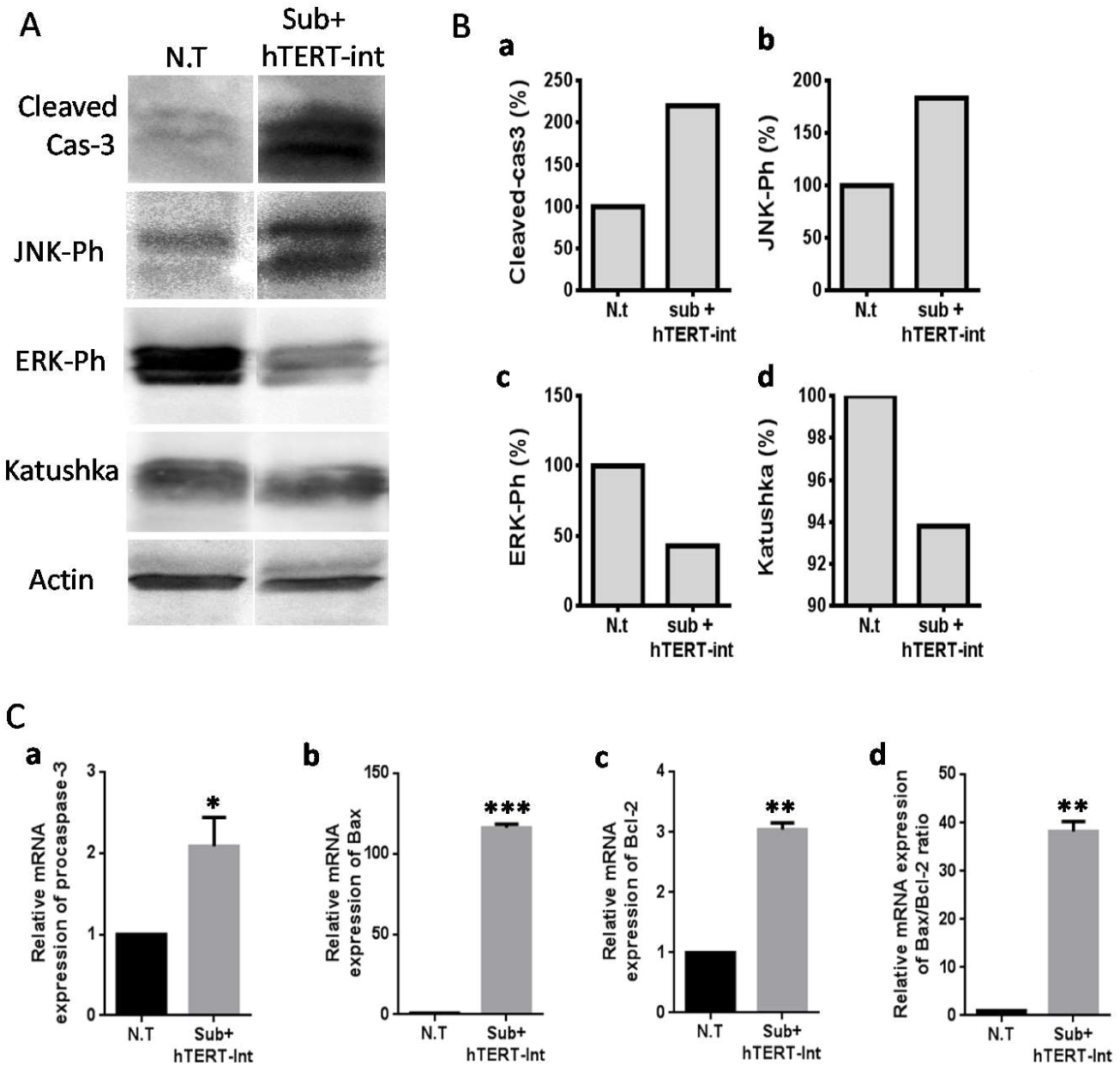
Figure 4. A. Western blot analysis of apoptosis-associated and proliferation proteins in LLC-Kat cancer lungs. The total lung cellular proteins extracted from LLC-Kat lungs were probed with Sub+*hTERT*-Int using antibodies recognizing the following apoptotic and proliferation markers: cleaved Cas-3, Ph-JNK, Ph-ERK, and Katushka. The actin marker was used as unrelated controls. N.T. – untreated lungs, Sub+*hTERT*-Int – the lungs treated with the Sub+ *hTERT*-Int plasmid. Each panel shows the typical lungs from a cohort of at least three mice. B. RT PCR analysis of apoptosis-associated genes in LLC-Kat cancer lungs treated with Sub+*hTERT*-Int. The relative mRNA expression levels normalized to Katushka were detected for (a) procaspase-3, (b) Bax, (c) Bcl-2, (d) Bax to Bcl-2 ratio. ***p-val ≤ 0.001; **p-val ≤ 0.01; *p-val ≤ 0.05 vs.N.T.

Since DTA expression induces apoptosis as described before (18;19), using qRT PCR we wanted to analyze the mini-transcriptome of the following genes: proCas-3, Bax that is functionally characterized as an apoptosis-promoting factor and the Bcl-2 that is characterized as an apoptosis-suppressing factor (52;53) in LLC-Kat cancer lungs treated by the binary system (Fig.4B, a, b, c, respectively). This analysis demonstrated that the level of Bax, Bcl-2 and proCas-3 gene transcription in cancer treated lungs were much higher than in the cancer untreated lungs (Fig.4B). Moreover, the qRT-PCR analysis also demonstrated an increased ratio of Bax/Bcl-2 in the LLC-Kat cancer lungs treated by the binary system compared to the untreated lungs. Bax/Bcl-2 ratio is described as a cellular ‘rheostat’ of apoptosis sensitivity in the sense that the intracellular ratio of bax/bcl-2 protein can profoundly influence the ability of a cell to respond to an apoptotic signal (54;55). According to this concept, a cell with a high bax/bcl-2 ratio will be more sensitive to a given apoptotic stimuli when compared to a similar cell type with a comparatively low bax/bcl-2 ratio (56). (Fig. 4B, d).

## DISCUSSION

In this paper, we have developed an anti-cancer binary system based on a site-specific recombination reaction catalyzed by Int that activates the expression of the dta toxin gene specifically in cancerous cells without affecting neighboring normal cell.

Comparative analyses of the Int-based binary and *hTERT-dta* mono systems in healthy C57BL/6 mice has revealed an unexpected substantial DTA expression in mice treated with *hTERT-dta* plasmid as a result of non-specific *hTERT* promoter leakage that led to the activation of apoptosis indicated by the apoptotic markers Cas-3 and P-53 and TUNEL, demonstrating the poor safety of the mono system [Fig. 2, panels (e)]. In contrast, the Int-based binary system did not cause the similar non-specific effect. These results demonstrate the efficacy and safety of the binary system towards normal vs. cancerous cells [Fig. 2, panels (e)]. As described above the comparative quantitative analysis of DTA expression level in LLC-Kat mice lungs Int-based binary system revealed elevation of 33 folds more DTA over the mono system (Fig. 3, e). Actually Int-catalyzed *attR* x *attL* recombination removes the terminator sequence from the silent substrate that produce the *hTERT-attB-dta* product plasmid which differs from *hTERT-dta* plasmid only the presence of the short (50 bp) *attB* sequence between the promoter and *dta* gene. We proposed that the difference in DTA expression in these two plasmids may be explained by *hTERT* promoter differential regulation in the binary and mono systems in LLC-Kat cancer cells.

*hTERT* promoter regulates the expression of human telomerase reverse transcriptase (*hTERT*) gene which product is a part of enzyme telomerase together with an RNA subunit and telomerase-associated proteins (57). Whereas the RNA subunit of telomerase is expressed in most cells, *hTERT* expression is repressed in the normal cells, while in more than 85% of all tumor cells *hTERT* transcription is upregulated. Thus, *hTERT* expression resulting in telomerase activity is critical for tumorigenesis (58). Furthermore, inhibition of telomerase activity leads to senescence or apoptosis in tumor cells (59-61), indicating that telomerase activity is required for the long-term viability of tumor cells. One of the most significant regulatory elements in *hTERT* promoter sequence are E boxes – the binding sites for the Myc/Max/Mad network of basic helixloop-helix/leucine zipper transcription factors with roles in various aspects of cell behavior including proliferation, differentiation and apoptosis (62). Central is Max that can both homodimerize and form heterodimers with Myc and Mad proteins, resulting in gene activation (Myc/Max) or repression (Mad/Max). The Mad family consists of four related protein designated Mad1, Mad2 (Mxi1), Mad3 and Mad4 and more distantly related members of the bHLH-ZIP family, Mnt and Mga. Like Myc, the Mad proteins are tightly regulated with short half-lives. In general, Mad family members interfere with Myc-mediated processes such as proliferation, transformation and prevention of apoptosis by inhibiting transcription (63-65). Thus, the antagonism between Myc and Mad proteins may be a crucial determinant of hTERT expression and telomerase activity (66).

We suggest that the appearance of a high concentration of active exogenous *hTERT* promoter provided by the *hTERT-dta* plasmid causes the sharp change of active *hTERT* intracellular balance that results in its immediate repression by the different transcription factors activation including Mad1. On the other hand, the *hTERT* promoter on the plasmid with the silent DTA substrate in the binary system is not active due to the intervention of the inserted transcription terminator. In this case the balance of active *hTERT* promoter in the cells remains invariable. Consequently, the silent dta substrate activation by Int-catalyzed site-specific recombination in LLC-Kat cancer cells does not lead to formation of the significant amount of *hTERT-attB-dta* product and as a result to an essential change of balance of active *hTERT* promoter. However, the amount of this product is suffice for the DTA synthesis in the cancer cell leading to the high efficacy killing activity as described in the results.

To test the proposed suggestion we have performed the analysis of the Mad1 expression level in both healthy and lung cancer mice by IHC using Ab against Mad1 (Fig. 5 A, B).

**Figure 5.**
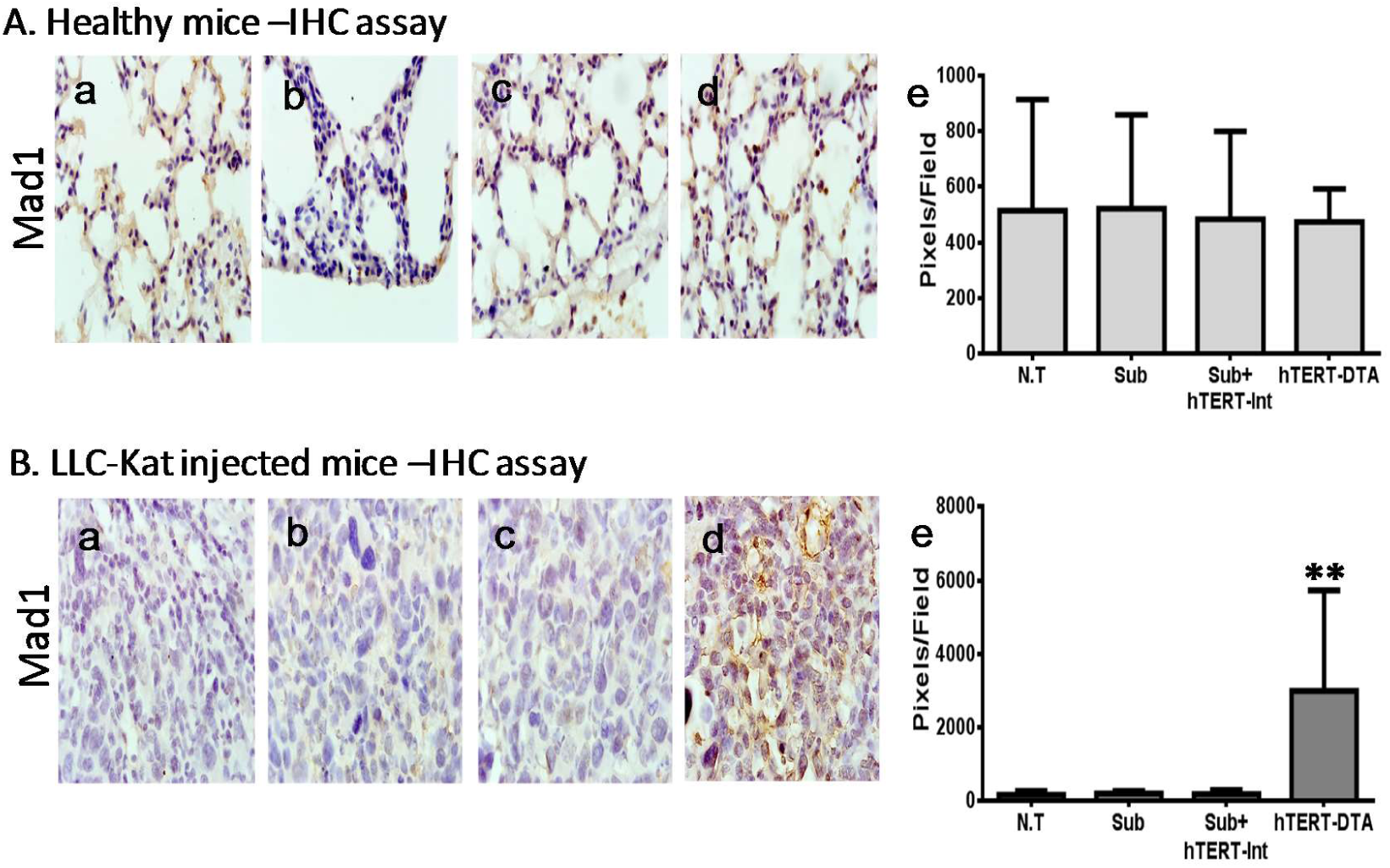
IHC analyses of Mad1 expression in the lungs of the healthy mice (A) and of the of LLC-Kat injected mice (B). The mice were treated in the same conditions as in Fig. 3. (a) Untreated mice (N.T.), (b) Sub, (c) Sub + *hTERT*-Int, (d) *hTERT*-DTA, (e) - quantitative data. The bars show the mean value of five experiments; the error bars indicate standard deviation. **p-val ≤ 0.01 vs. N.T.

As expected, Fig. 5, A shows that in healthy lungs, the silent dta substrate, the binary system and the *hTERT- dta* plasmid [panels (b, c and d) respectively] did not show any immune detection as in the untreated group [panels (a)].

Intriguingly, among all four LLC-Kat lung cancer mice groups only the group treated with *hTERT-dta* plasmid showed Mad1 positive immune detection [panels (d)] justifying the regulatory interfering hypothesis: the exogenous active *hTERT* promoter repression occurs as a result of the Mad1 transcription factor elevation [Fig. 5, B, panels(d, e)]. Comparative quantitative analysis of Mad1 in LLC-Kat mice lungs between Int-based binary and the mono systems has revealed 33 fold increasing Mad1 expression level in the lungs treated with *hTERT- dta* plasmid [Fig. 5, B, panel (e)]. As shown in the results the comparative quantitative analysis of DTA expression level in healthy and LLC-Kat mice lungs treated with *hTERT-dta* plasmid has revealed an unexpected 11 folds increasing DTA level in healthy mice (Figs. 2, 3, e).This unforeseen result can be explained by the Mad1 down regulation effect. Indeed, the comparative quantitative analysis of Mad1 expression level in healthy and LLC-Kat mice lungs treated with *hTERT-dta* plasmid has revealed six folds increasing Mad1 level in LLC-Kat mice [Fig. 5, A, B, panels(e)] that led to more efficient down regulation of *hTERT* and as a result to more efficient DTA synthesis repression.

These data unambiguously demonstrates our suggestion that active *hTERT* intracellular down regulation is caused by activation and elevation of the suppressor regulator Mad1, different transcription factors that regulates *hTERT* promoter might be involved.

One of the Int distinctiveness that can significantly increase specificity and efficiency of the binary system in human cells is Int activity dependence on the host encoded integration host factor (IHF) and the phage-encoded excisionase (Xis) in E. coli (30).

We have shown that in contrast to the requirement of Int for IHF and Xis in its natural milieu, the wild type Int was active in human cells without the need to supply any accessory proteins (67). However the addition of these proteins improved the Int activity in human cells (68). Similar results have been also received for Int Lambda (Int-λ) in human cells. The expression of single chain IHF strengthened the wild type Int-λ activity to the levels of the IHF-independent mutants (69;70). In vitro experiments with Int-λ have also showed that human chromatin-associated proteins HMG1 and HMG2 can substitute to some extent for the requirement of the accessory proteins (71).

The results stated above allow assuming that definite histone, histone-like or chromatin-associated proteins can render the strengthening effect on Int activity in human cells. Intriguingly the expression of some human histones and histone-like proteins (H2A, H2B, H3, H4, CENP-A) are upregulated in numerous cancers (72-74). All these data allow suggesting that Int activity may be strengthened in the human cancer cells compare to healthy cells by the putative histone or histone-like Int accessory proteins upregulating. This phenomenon of Int cancer specific activation via such accessory proteins can significantly increase specificity and efficiency of the Int-based binary system that provides its additional advantage compare to conventional mono systems for human cancer therapy. For further cancer gene therapy studies we suggest to consider the regulation elements as a crucial and significant factor for the activity and efficiency of the developed approaches.

Our binary anti-cancer system belongs to the toxin gene delivery approach but has the obvious advantages compared to conventional toxin delivery approaches: it shows a higher safety efficacy in healthy cells and a higher killing efficacy in cancer cells. Moreover, unlike other systems using DNA vectors for mammalian gene manipulations ours is viral-free. This binary system may have a strong impact on human cancer curing. The preclinical trials including the examining of the binary system in survival experiments are in progress.

## ACKNOWLEGMENTS

We thank A. Shtulkarc for pKS1161 plasmids construction, M. Athamna for help with Real-Time PCR technique. This work was supported by the Kamin fund of the Israel Ministry of Economy and Industry. A.E. is thankful also for the support of the Constantiner Institute.

## Author Contributions

A.E., I.S., N.G., R. G. and M.K. designed research. A.E., I.S., N.G., R.G. and Y.Z. performed experiments. A.E., I.S., N.G., Y.Z., R.G. and M.K. analyzed data. A.E., M.K., R.G. and G.P. wrote the paper.

## Reference List

1. Santiago-Ortiz, J.L. and Schaffer, D.V. (2016) Adeno-associated virus (AAV) vectors in cancer gene therapy. J. Control Release, 240, 287–301.

2. Lattime E. and Gerson S. eds., Gene Therapy of Cancer: Translational Approaches from Preclinical Studies to Clinical Implementation. (2014) Academic press.

3. Everett, W.H. and Curiel, D.T. (2015) Gene therapy for radioprotection. Cancer Gene Ther., 22, 172–180.

4. Mayer, E.L. (2013) Early and late long-term effects of adjuvant chemotherapy. Am. Soc. Clin. Oncol. Educ. Book., 9–14.

5. Persidis, A. (1999) Cancer multidrug resistance. Nat. Biotechnol., 17, 94–95.

6. Nagengast, W.B., Oude Munnink, T.H., Dijkers, E.C., Hospers, G.A., Brouwers, A.H., Schroder, C.P., Lub-de, H.M. and de Vries, E.G. (2010) Multidrug resistance in oncology and beyond: from imaging of drug efflux pumps to cellular drug targets. Methods Mol. Biol., 596, 15–31.

7. Sharma, P. and Allison, J.P. (2015) The future of immune checkpoint therapy. Science, 348, 56–61.

8. Baek, S., Kim, C.S., Kim, S.B., Kim, Y.M., Kwon, S.W., Kim, Y., Kim, H. and Lee, H. (2011) Combination therapy of renal cell carcinoma or breast cancer patients with dendritic cell vaccine and IL-2: results from a phase I/II trial. J. Transl. Med., 9, 178.

9. Oldfield, E.H., Ram, Z., Culver, K.W., Blaese, R.M., DeVroom, H.L. and Anderson, W.F. (1993) Gene therapy for the treatment of brain tumors using intra-tumoral transduction with the thymidine kinase gene and intravenous ganciclovir. Hum. Gene Ther., 4, 39–69.

10. Maeda, H., Wu, J., Sawa, T., Matsumura, Y. and Hori, K. (2000) Tumor vascular permeability and the EPR effect in macromolecular therapeutics: a review. J. Control Release, 65, 271–284.

11. Maeda, H. (2001) The enhanced permeability and retention (EPR) effect in tumor vasculature: the key role of tumor-selective macromolecular drug targeting. Adv. Enzyme Regul., 41, 189–207.

12. Byrne, J.D., Betancourt, T. and Brannon-Peppas, L. (2008) Active targeting schemes for nanoparticle systems in cancer therapeutics. Adv. Drug Deliv. Rev., 60, 1615–1626.

13. Lee, H.Y., Mohammed, K.A. and Nasreen, N. (2016) Nanoparticle-based targeted gene therapy for lung cancer. Am. J. Cancer Res., 6, 1118–1134.

14. Showalter, S.L., Huang, Y.H., Witkiewicz, A., Costantino, C.L., Yeo, C.J., Green, J.J., Langer, R., Anderson, D.G., Sawicki, J.A. and Brody, J.R. (2008) Nanoparticulate delivery of diphtheria toxin DNA effectively kills Mesothelin expressing pancreatic cancer cells. Cancer Biol. Ther., 7, 1584–1590.

15. Amit, D., Tamir, S. and Hochberg, A. (2013) Development of targeted therapy for a broad spectrum of solid tumors mediated by a double promoter plasmid expressing diphtheria toxin under the control of IGF2-P4 and IGF2-P3 regulatory sequences. Int. J. Clin. Exp. Med., 6, 110–118.

16. Cocco, E., Deng, Y., Shapiro, E.M., Bortolomai, I., Lopez, S., Lin, K., Bellone, S., Cui, J., Menderes, G., Black, J.D. et al. (2017) Dual-Targeting Nanoparticles for In Vivo Delivery of Suicide Genes to Chemotherapy-Resistant Ovarian Cancer Cells. Mol. Cancer Ther., 16, 323–333.

17. Collier, R.J. (2001) Understanding the mode of action of diphtheria toxin: a perspective on progress during the 20th century. Toxicon, 39, 1793–1803.

18. Perentesis, J.P., Waddick, K.G., Bendel, A.E., Shao, Y., Warman, B.E., Chandan-Langlie, M. and Uckun, F.M. (1997) Induction of apoptosis in multidrug-resistant and radiation-resistant acute myeloid leukemia cells by a recombinant fusion toxin directed against the human granulocyte macrophage colony-stimulating factor receptor. Clin. Cancer Res., 3, 347–355.

19. Thorburn, J., Frankel, A.E. and Thorburn, A. (2003) Apoptosis by leukemia cell-targeted diphtheria toxin occurs via receptor-independent activation of Fas-associated death domain protein. Clin. Cancer Res., 9, 861–865.

20. Lee, E.J. and Jameson, J.L. (2002) Cell-specific Cre-mediated activation of the diphtheria toxin gene in pituitary tumor cells: potential for cytotoxic gene therapy. Hum. Gene Ther., 13, 533–542.

21. Massie, B., Couture, F., Lamoureux, L., Mosser, D.D., Guilbault, C., Jolicoeur, P., Belanger, F. and Langelier, Y. (1998) Inducible overexpression of a toxic protein by an adenovirus vector with a tetracycline-regulatable expression cassette. J. Virol., 72, 2289–2296.

22. Branda, C.S. and Dymecki, S.M. (2004) Talking about a revolution: The impact of site-specific recombinases on genetic analyses in mice. Developmental Cell, 6, 7–28.

23. Glaser, S., Anastassiadis, K. and Stewart, A.F. (2005) Current issues in mouse genome engineering. Nature Genetics, 37, 1187–1193.

24. Wirth, D., Gama-Norton, L., Riemer, P., Sandhu, U., Schucht, R. and Hauser, H. (2007) Road to precision: recombinase-based targeting technologies for genome engineering. Curr. Opin. Biothecnol., 18, 411–419.

25. Nafissi, N. and Slavcev, R. (2014) Bacteriophage recombination systems and biotechnical applications. Appl. Microbiol. Biotechnol., 98, 2841–2851.

26. Krappmann, S. (2014) Genetic surgery in fungi: employing site-specific recombinases for genome manipulation. Appl. Microbiol. Biotechnol., 98, 1971–1982.

27. Gaj, T., Sirk, S.J. and Barbas, C.F., III (2014) Expanding the scope of site-specific recombinases for genetic and metabolic engineering. Biotechnol. Bioeng., 111, 1–15.

28. Grindley, N., Whitestone, K. and Rice, P. (2006) Mechanism of site-specific recombination. Annu. Rev. Biochem., 75, 567–605.

29. Azaro, M.A. and Landy, A. (2002) Integrase and the l int family. In Craig, N.L., Craigie, R., Gellert, M. and Lambowitz, A. (eds.), Mobile DNAII. ASM Press, Washington DC, pp. 118–148.

30. Weisberg, R.A., Gottesmann, M.E., Hendrix, R.W. and Little, J.W. (1999) Family values in the age of genomics: comparative analyses of temperate bacteriophage HK022. Annu. Rev. Genet., 33, 565–602.

31. Gottfried, P., Lotan, O., Kolot, M., Maslenin, L., Bendov R, Gorovits, R., Yesodi, V., Yagil, E. and Rosner, A. (2005) Site-specific recombination in Arabidopsis plants promoted by the Integrase protein of coliphage HK022. Plant Molecular Biology, 57, 435–444.

32. Voziyanova, E., Malchin, N., Anderson, R.P., Yagil, E., Kolot, M. and Voziyanov, Y. (2013) Efficient Flp-Int HK022 dual RMCE in mammalian cells. Nucleic Acids Res., 41, e125.

33. Elias, A., Spector, I., Sogolovsky-Bard, I., Gritsenko, N., Rask, L., Mainbakh, Y., Zilberstein, Y., Yagil, E. and Kolot, M. (2016) Cancer-specific binary expression system activated in mice by bacteriophage HK022 Integrase. Sci. Rep., 6, 24971.

34. Dorgai, L., Yagil, E. and Weisberg, R. (1995) Identifying determinants of recombination specificity: construction and characterization of mutant bacteriophage integrases. J. Mol. Biol., 252, 178–188.

35. Sambrook, J., Fritsch, E.F. and Maniatis, T. (1989) Molecular cloning: a laboratory manual. Cold Spring Harbor Laboratory, Cold Spring Harbor, NY.

36. Rask, L., Fregil, M., Hogdall, E., Mitchelmore, C. and Eriksen, J. (2013) Development of a metastatic fluorescent Lewis Lung carcinoma mouse model: identification of mRNAs and microRNAs involved in tumor invasion. Gene, 517, 72–81.

37. Maxwell, F., Maxwell, I.H. and Glode, L.M. (1987) Cloning, sequence determination, and expression in transfected cells of the coding sequence for the tox 176 attenuated diphtheria toxin A chain. Mol. Cell Biol., 7, 1576–1579.

38. Cong, Y.S., Wen, J. and Bacchetti, S. (1999) The human telomerase catalytic subunit *hTERT*: organization of the gene and characterization of the promoter. Hum. Mol. Genet., 8, 137–142.

39. Majumdar, A.S., Hughes, D.E., Lichtsteiner, S.P., Wang, Z., Lebkowski, J.S. and Vasserot, A.P. (2001) The telomerase reverse transcriptase promoter drives efficacious tumor suicide gene therapy while preventing hepatotoxicity encountered with constitutive promoters. Gene Ther., 8, 568–578.

40. Sauer, B. (1993) Manipulation of transgenes by site-specific recombination: use of Cre recombinase. Methods Enzymol., 225, 890–900.

41. Counter, C.M., Avilion, A.A., LeFeuvre, C.E., Stewart, N.G., Greider, C.W., Harley, C.B. and Bacchetti, S. (1992) Telomere shortening associated with chromosome instability is arrested in immortal cells which express telomerase activity. EMBO J., 11, 1921–1929.

42. Xi, L., Schmidt, J.C., Zaug, A.J., Ascarrunz, D.R. and Cech, T.R. (2015) A novel two-step genome editing strategy with CRISPR-Cas9 provides new insights into telomerase action and TERT gene expression. Genome Biol., 16, 231.

43. Bertram, J.S. and Janik, P. (1980) Establishment of a cloned line of Lewis Lung Carcinoma cells adapted to cell culture. Cancer Lett., 11, 63–73.

44. Shcherbo, D., Merzlyak, E.M., Chepurnykh, T.V., Fradkov, A.F., Ermakova, G.V., Solovieva, E.A., Lukyanov, K.A., Bogdanova, E.A., Zaraisky, A.G., Lukyanov, S. et al. (2007) Bright far-red fluorescent protein for whole-body imaging. Nat. Methods, 4, 741–746.

45. Elmore, S. (2007) Apoptosis: a review of programmed cell death. Toxicol. Pathol., 35, 495–516.

46. Fridman, J.S. and Lowe, S.W. (2003) Control of apoptosis by p53. Oncogene, 22, 9030–9040.

47. Weigel, M.T., Rath, K., Alkatout, I., Wenners, A.S., Schem, C., Maass, N., Jonat, W. and Mundhenke, C. (2014) Nilotinib in combination with carboplatin and paclitaxel is a candidate for ovarian cancer treatment. Oncology, 87, 232–245.

48. Behrens, A., Sabapathy, K., Graef, I., Cleary, M., Crabtree, G.R. and Wagner, E.F. (2001) Jun N-terminal kinase 2 modulates thymocyte apoptosis and T cell activation through c-Jun and nuclear factor of activated T cell (NF-AT). Proc. Natl. Acad. Sci. U. S. A, 98, 1769–1774.

49. Makena, P.S., Gorantla, V.K., Ghosh, M.C., Bezawada, L., Balazs, L., Luellen, C., Parthasarathi, K., Waters, C.M. and Sinclair, S.E. (2011) Lung injury caused by high tidal volume mechanical ventilation and hyperoxia is dependent on oxidant-mediated c-Jun NH2-terminal kinase activation. J. Appl. Physiol (1985.), 111, 1467–1476.

50. Xia, Y., Yeddula, N., Leblanc, M., Ke, E., Zhang, Y., Oldfield, E., Shaw, R.J. and Verma, I.M. (2012) Reduced cell proliferation by IKK2 depletion in a mouse lung-cancer model. Nat. Cell Biol., 14, 257–265.

51. Yu, Y., Luk, F., Yang, J.L. and Walsh, W.R. (2011) Ras/Raf/MEK/ERK pathway is associated with lung metastasis of osteosarcoma in an orthotopic mouse model. Anticancer Res., 31, 1147–1152.

52. Hata, A.N., Engelman, J.A. and Faber, A.C. (2015) The BCL2 Family: Key Mediators of the Apoptotic Response to Targeted Anticancer Therapeutics. Cancer Discov., 5, 475–487.

53. Adams, J.M. and Cory, S. (2007) Bcl-2-regulated apoptosis: mechanism and therapeutic potential. Curr. Opin. Immunol., 19, 488–496.

54. Yang, E. and Korsmeyer, S.J. (1996) Molecular thanatopsis: a discourse on the BCL2 family and cell death. Blood, 88, 386–401.

55. Oltvai, Z.N., Milliman, C.L. and Korsmeyer, S.J. (1993) Bcl-2 heterodimerizes *in vivo* with a conserved homolog, Bax, that accelerates programmed cell death. Cell, 74, 609–619.

56. Perlman, H., Zhang, X., Chen, M.W., Walsh, K. and Buttyan, R. (1999) An elevated bax/bcl-2 ratio corresponds with the onset of prostate epithelial cell apoptosis. Cell Death. Differ., 6, 48–54.

57. Nugent, C.I. and Lundblad, V. (1998) The telomerase reverse transcriptase: components and regulation. Genes Dev., 12, 1073–1085.

58. Hiyama, E. and Hiyama, K. (2003) Telomerase as tumor marker. Cancer Lett., 194, 221–233.

59. Hahn, W.C., Stewart, S.A., Brooks, M.W., York, S.G., Eaton, E., Kurachi, A., Beijersbergen, R.L., Knoll, J.H., Meyerson, M. and Weinberg, R.A. (1999) Inhibition of telomerase limits the growth of human cancer cells. Nat. Med., 5, 1164–1170.

60. Herbert, B., Pitts, A.E., Baker, S.I., Hamilton, S.E., Wright, W.E., Shay, J.W. and Corey, D.R. (1999) Inhibition of human telomerase in immortal human cells leads to progressive telomere shortening and cell death. Proc. Natl. Acad. Sci. U. S. A, 96, 14276–14281.

61. Zhang, X., Mar, V., Zhou, W., Harrington, L. and Robinson, M.O. (1999) Telomere shortening and apoptosis in telomerase-inhibited human tumor cells. Genes Dev., 13, 2388–2399.

62. Baudino, T.A. and Cleveland, J.L. (2001) The Max network gone mad. Mol. Cell Biol., 21, 691–702.

63. Henriksson, M. and Luscher, B. (1996) Proteins of the Myc network: essential regulators of cell growth and differentiation. Adv. Cancer Res., 68, 109–182.

64. Grandori, C., Cowley, S.M., James, L.P. and Eisenman, R.N. (2000) The Myc/Max/Mad network and the transcriptional control of cell behavior. Annu. Rev. Cell Dev. Biol., 16, 653–699.

65. Luscher, B. (2012) MAD1 and its life as a MYC antagonist: an update. Eur. J. Cell Biol., 91, 506–514.

66. Janknecht, R. (2004) On the road to immortality: *hTERT* upregulation in cancer cells. FEBS Lett., 564, 9–13.

67. Harel-Levy G., Goltsman J., Tuby C.N.J.H., Yagil E. and Kolot, M. (2008) Human genomic site-specific recombination catalyzed by coliphge HK022 integrase. J. Biotechnol., 134, 45–54.

68. Malchin, N., Goltsman, J., Dabool, L., Gorovits, R., Bao, Q., Droge, P., Yagil, E. and Kolot, M. (2009) Optimization of coliphage HK022 Integrase activity in human cells. Gene, 437, 9–13.

69. Christ, N., Corona, T. and Droge, P. (2002) Site-specific recombination in eukaryotic cells mediated by mutant lambda integrases: Implications for synaptic complex formation and the reactivity of episomal DNA segments. J. Mol. Biol., 319, 305–314.

70. Corona, T., Bao, Q.Y., Christ, N., Schwartz, T., Li, J.M. and Droge, P. (2003) Activation of site-specific DNA integration in human cells by a single chain integration host factor. Nucleic Acids Research, 31, 5140–5148.

71. Segall, A.M., Goodman, S.D. and Nash, H.A. (1994) Architectural elements in nucleoprotein complexes: interchangeability of specific and non-specific DNA binding proteins. EMBO J., 13, 4536–4548.

72. Gao, J., Aksoy, B.A., Dogrusoz, U., Dresdner, G., Gross, B., Sumer, S.O., Sun, Y., Jacobsen, A., Sinha, R., Larsson, E. et al. (2013) Integrative analysis of complex cancer genomics and clinical profiles using the cBioPortal. Sci. Signal., 6, l1.

73. Cerami, E., Gao, J., Dogrusoz, U., Gross, B.E., Sumer, S.O., Aksoy, B.A., Jacobsen, A., Byrne, C.J., Heuer, M.L., Larsson, E. et al. (2012) The cBio cancer genomics portal: an open platform for exploring multidimensional cancer genomics data. Cancer Discov., 2, 401–404.

74. Maze, I., Noh, K.M., Soshnev, A.A. and Allis, C.D. (2014) Every amino acid matters: essential contributions of histone variants to mammalian development and disease. Nat. Rev. Genet., 15, 259–271.

75. Kolot, M., Meroz, A. and Yagil, E. (2003) Site-specific recombination in human cells catalyzed by the wild-type integrase protein of coliphage HK022. Biotechnol. Bioeng., 84, 56–60.

